# Protective human IgE responses are promoted by comparable life-cycle dependent Tegument Allergen-Like protein expression in *Schistosoma haematobium* and *Schistosoma mansoni* infection

**DOI:** 10.1101/2022.12.02.518813

**Authors:** Rebecca C. Oettle, Harriet A. Dickinson, Colin M. Fitzsimmons, Moussa Sacko, Edridah M. Tukahebwa, Iain W. Chalmers, Shona Wilson

## Abstract

*Schistosoma haematobium* is the most prevalent of the human-infecting schistosome species, causing significant morbidity in endemically exposed populations. Despite this, it has been relatively understudied compared to its fellow species, *S. mansoni*. Here we provide the first comprehensive characterization of the *S. haematobium* Tegument Allergen-Like protein family, a key protein family directly linked to protective immunity in *S. mansoni* infection. Comparable with observations for *S. mansoni*, parasite phylogenetic analysis and relative gene expression combined with host serological analysis support a cross-reactive relationship between *S. haematobium* TAL proteins, exposed to the host immune system as adult worms die, and closely related proteins, exposed during penetration by the infecting cercarial and early schistosomulae stages. Specifically, our results strengthen the evidence for host immunity driven by cross-reactivity between family members TAL3 and TAL5, establishing it for the first time for *S. haematobium* infection. Furthermore, we build upon this relationship to include the involvement of an additional member of the TAL protein family, TAL11 for both schistosome species. Finally, we show a close association between experience of infection and intensity of transmission and the development of protective IgE responses to these antigens, thus improving our knowledge of the mechanisms by which protective host immune responses develop. This knowledge will be critical in understanding how control efforts such as mass drug administration campaigns influence the development of host immunity and subsequent patterns of infection and disease within endemic populations.

**Author Summary:** *S. haematobium* is the most prevalent of the human infecting schistosomes. Along with *S. mansoni*, it is responsible for the majority of schistosomiasis cases that are borne by the populations of sub-Saharan Africa, where the global burden of this infection is centered. Here, we provide insight into the IgE antibody response that protects against these infections. Through utilization of *in silico* analysis and transcriptional studies of parasite life stages, in combination with immuno-epidemiological studies, we explore the relationship between host immune protection and a parasite protein family named the Tegument Allergen-Like (TAL) proteins. Our results show that several members of the TAL protein family are important in host protection to both these major schistosome species. For the first time we demonstrate that a progressive cross-reactive TAL-IgE response occurs against *S. haematobium*, similar to that previous observed in *S. mansoni* infection. We additionally expand upon previous knowledge for *S. mansoni*, identifying further complexity in the cross-reactive relationship between TAL family members, providing evidence of a key role for family member TAL11 in induction of the protective host immune response.

## Background

Schistosomiasis is a disease of considerable public health importance resulting in severe morbidity and reduced quality of life in over 290 million people affected by the disease worldwide (1). The major burden of infection is held within sub-Saharan Africa where *Schistosoma haematobium* and *S. mansoni* are the prevalent infections. The acquisition of protective immunity against infection with *Schistosoma* worms is a gradual process thought to naturally occur over several years, however, the mechanism is not fully understood. Evidence from immuno-epidemiological studies principally supports the hypothesis that a humoral response is acquired following cumulative exposure to antigens that are released as adult worms die (2). IgE is the main class of protective antibody (4,5) and in a process named delayed-concomitant immunity, the invading cercarial stage and the subsequent early schistosomule stage of infection are the proposed targets of the protective response (3). Whilst IgE stimulating antigens are not considered appropriate vaccine candidates, due the association between IgE and anaphylactic responses (6), improving our knowledge of the protective response, and its development in relationship to accumulative infection exposure, is essential for understanding *Schistosoma* transmission dynamics and subsequent planning of infection control measures.

In *S. mansoni* infection, members of a parasite protein family, the Tegument Allergen-Like (TAL) proteins, induce IgE responses that are strongly associated with human immunity (7,8). The TAL proteins form a family of calcium binding proteins unique to the platyhelminths (11) and proteins within the TAL family have been characterized in several *Schistosoma* species (5,9,10,12,13) in addition to other parasitic flatworms, including *Fasciola hepatica, F. gigantica, Clonorchis sinensis* and *Opisthorchis viverrini* (14–17). The family have high levels of sequence and predicted structural similarity, typically consisting of a N-terminal domain containing two EF-hand-like structures (InterPro domain: IPR011992) and a C-terminal dynein light chain (DLC) domain (InterPro domain: IPR037177, IPR001372) (8,11). EF-hand motifs are a known feature of many IgE antigens in both plants and animals (18) and comprise one of the largest allergenic protein domain families (19). The first members of the TAL family to be independently identified in *S. mansoni* were SmTAL1 (5), SmTAL2 (21), and SmTAL3 (22), previously named Sm22.6, Sm21.7 and Sm20.8, respectively. These proteins have since been characterized in depth (8,11), along with more recent molecular and biochemical characterization of ten additional members of the family (SmTAL4-13) (8,12).

Several studies have provided evidence for strong cross-reactivity amongst IgE responses to EF-hand containing proteins, resulting in cross-sensitization between environmental and food allergens (23–25) and it has been proposed that members of the TAL protein family may stimulate the protective response to schistosomes via cross-reactive antigen recognition (3). Indeed, immune cross-reactivity has recently been established for another family of schistosome proteins, the Venom Allergen-Like proteins (26). mRNA expression data shows that the TAL proteins are expressed at varying levels across the different life cycle stages in S. *mansoni* (8). Of interest regarding cross-reactive immunity, SmTAL1, SmTAL3 and SmTAL11 are largely expressed in the adult worm, whereas SmTAL4 and SmTAL5 are predominantly expressed in the cercaria (8). Whole-mount immunostaining of cercarial sections demonstrates that SmTAL4 expression is restricted to the tail of the cercaria (8), which separates from the cercarial head during skin penetration. SmTAL4 is therefore not indicated as an immune stimulatory protein or a target of the protective response. Conversely, SmTAL5 is expressed throughout the cercaria and early schistosomula stages and is therefore likely exposed to the immune system during invasion and early transformation. Regarding the TAL proteins expressed in the adult worm, SmTAL1 is the dominant antigen detected by IgE in immune sera. In endemic areas, the response increases in populations with age and corresponds with a decrease in infection intensity (8). However, IgE to SmTAL1 does not exhibit cross-reactivity with the cercarial expressed SmTAL5, whilst SmTAL3 does. SmTAL3, rather than SmTAL5, appears to be the antigenic source for this cross-reactive IgE response (3) and it is this relationship that is proposed to drive delayed concomitant immunity. The IgE response to the remaining adult worm expressed protein, SmTAL11, has not been characterized.

To date, only one member of the *S. haematobium* TAL protein family has been identified, a TAL1 orthologue which has a closely related sequence to the corresponding *S. mansoni* (9). Furthermore, the relationship between TAL-specific antibodies and the acquisition of protective immunity is yet to be defined in *S. haematobium*, although there is evidence for similar TAL1-specific IgE responses to those observed for *S. mansoni* (9, 27). Therefore, following publication of the *S. haematobium* genome and the subsequent updated version (28), we take the opportunity to characterize the *S. haematobium* TAL proteins, specifically those proteins orthologous to the *S. mansoni* proteins associated with protective IgE responses: ShTAL1, ShTAL3, and ShTAL5. We also take the opportunity to further characterize the *S. mansoni* protective response by testing the hypothesis that SmTAL11 induces an IgE response following worm death that contributes to the development of delayed concomitant immunity, and to examine the corresponding response in *S. haematobium* infection.

## Materials and methods

### Study population and sampling

IgE responses to the TAL protein family were examined across two separate study populations: one a fishing community situated on Lake Victoria, endemic for *S. mansoni;* and the other from three villages in the Segou Region, Mali, located along the River Niger and one of its tributaries, that are endemic for *S. haematobium*. Full descriptions of the study cohorts can be found elsewhere (3, 29). Quantitative parasitology was conducted for each individual at baseline prior to treatment, 9-weeks post-treatment with 40 mg/kg praziquantel and again at two years post-treatment. For the *S. mansoni* endemic Ugandan cohort this comprised Kato Katz egg counts performed in duplicate on three stool samples collected on different days. Similarly, for the *S. haematobium* endemic Malian cohort, egg counts were performed on three 10 ml filtered urine samples, collected on different days. Blood samples were collected into EDTA from both cohorts at baseline and 9-weeks post-treatment and the plasma harvested by centrifugation. Plasma samples were treated with 0.3 % tributyl phosphate/1 % Tween 80 to inactivate encapsulated viruses prior to measurement of specific antibody responses. CAA measurement was performed on a 40 μl sample of plasma from each participant in the Malian cohort at Leiden University Medical Center (LUMC) as previously described (30).

### Ethics statement

Ethical clearance was granted by the Uganda National Council of Science and Technology (Reference number: UNCST: HS59) and the Malian Ethical Review Committee of the National Institute for Research in Public Health (decision 0002/INRSP/DAP/SP-2005), respectively. In Uganda, consent forms were translated into the local language and informed written consent was obtained from all adults and from the parents/legal guardians of all children under 15. Whilst, in Mali, oral informed consent was given by adult participants and by parents or guardians of participating children. At the time of the study oral consent was deemed acceptable by the Malian Ministry of Health, due to cultural reasons and low literacy rates in the Malian villages.

### Parasite materials

*S. mansoni* parasite material was a gift from Prof David Dunne (University of Cambridge). Total RNA was isolated as previously described (31). *S. haematobium* material was kindly provided by Prof Ron Hokke (Leiden University Medical Center, Netherlands) and processed as described for *S. mansoni*.

### Bioinformatic Analysis

Amino acid sequences from well-characterized *S. mansoni* TAL proteins (8,11,12) were used in BLASTp searches against the predicted protein database of the recently published *S. haematobium* genome assembly (SchHae_3.0; NCBI BioProject: PRJNA78265; (28)). In line with the SmTAL standards reported in Fitzsimmons *et al*. (8) the criteria for ShTAL identification were: 1) overall sequence similarity (E<1^-10^); 2) the presence of at least one predicted N-terminal EF-hand domain (InterPro domain: IPR011992), and 3) a C-terminal dynein light chain (DLC) domain (InterPro domain: IPR037177, IPR001372), with an exception for TAL10, for which the *S. mansoni* orthologue was previously found to contain a DLC region that is not predicted by domain predication tools (8). Where no significant orthologous protein was identified, any remaining sequences that were unassigned to an orthologous *S. mansoni* TAL were used in BLASTp searches against the predicted protein database of the *S. mansoni* genome assembly (version 9, GCA_000237925.3).

### Alignment and Phylogenetic trees

Sequences were aligned using the Multiple Sequence Comparison by Log-Expectation tool (version 3.8 (32)) and conserved domains identified using Gblocks software (33), using the following conservative parameters: conserved or flanking positions were identified in a minimum of 50 % of the submitted sequences; blocks had a minimum length of five amino acids; positions with a gap in fewer than 50 % of the sequences could be selected in the final alignment, and the maximum number of contiguous non-conserved positions was limited to 8. The alignment was visualized in in Jalview (Version 2.103b (34)). A phylogenetic tree of the conserved regions was subsequently constructed in MEGAII (35) using a Maximum Likelihood method based on the JTT matrix-based model (36). Bootstrapping was performed over 1000 iterations with fewer than 20 % alignment gaps, missing data or ambiguous bases allowed at any position. All positions with less than 50 % site coverage were eliminated. Following analysis, the Gblock output dataset contained a total of 101 conserved positions.

### Real-time PCR analysis

The transcript abundance of sequence verified *Shtal1, Shtal3, Shtal5, Shtal11, Smtal1, Smtal3, Smtal5*, and *Smtal11* mRNA was quantified relative to Long Ribosomal Protein II (RPII) (Sha_102687; Shaem_V3.0 locus tag MS3_00007736; Smp_156670) in egg, cercariae and adult worm stages of the *S. haematobium* lifecycle. RPII was selected as a reference gene in the qPCR analysis from a list of suggested housekeeping genes that demonstrate stable expression across four lifecycle stages, following analysis by NormFinder (37). Furthermore, RPII homologues can be identified across the *Schistosoma* species (38–40). The cDNA from each life cycle stage was assayed by real-time (RT) PCR using gene-specific primers (Supporting Information Table S1). The amplification efficiency of each primer pair was determined by plotting cycle thresholds from serial two-fold dilutions of suitable cDNA samples to form a five-point standard curve. Control reactions lacking the cDNA template were run for each primer set and a no primer control was included on each plate, to confirm the absence of non-specific amplification in PCR reactions. An additional ‘mix’ stage control, comprising the three lifecycle stages in equal proportions was included as a reference control. All reactions were performed in triplicate on an Eco Real-Time PCR machine (Illumina) using KAPA SYBER FAST (KAPA Biosciences), according to the manufacturer’s instructions. Relative quantification was performed using the delta-delta Ct, or Livak method (41), to normalize expression of the target genes relative to the reference gene (RPII) and relative to the reference ‘mix’ sample. The following equation was used to calculate a normalized ShTAL expression ratio using the Illumina Eco system software (Illumina):

Fold gene expression = 2^-(ΔΔCt), where ΔΔCt =ΔCt (Sample) – ΔCt (Control average).

The resulting data were compared to transcription analysis published alongside the *S. haematobium* version 3 genome annotation (28).

### Recombinant protein production

The full coding transcript sequences were cloned from cDNA from adult *S. haematobium* worms, Egyptian Strain, ligated into the expression vector pGEX-1λT (Pharmacia) and expressed as a fusion protein with glutathione-S-transferase (GST), as described previously (42). Recombinant plasmids were then isolated (MiniPrep, Qiagen) and sequenced (DNA Sequencing Facility, University of Cambridge). Recombinant proteins were expressed and purified as described previously (8).

### Human IgE ELISA and reciprocal inhibition assay

Human IgE and IgG4 ELISAs were performed to measure the TAL antigen specific IgE and IgG4 antibody concentration in individual plasma samples from each study site. The saturating coating concentration for each recombinant antigen was identified using a coating inhibition assay, as described elsewhere (8) and ELISA plates were coated with the determined recombinant TAL antigen concentration: between 3 μg/ml and 9 μg/ml (Supporting Information Table S2). A serial dilution of commercial, purified human immunoglobulin protein was added to each plate to provide a reference standard curve, IgE (Calbiochem) and IgG4 (Sigma). Plates were incubated overnight at 4°C, washed, and blocked with 1 % Marvel milk powder in 1 x PBS for 1 hour. Plasma, diluted 1:20 for IgE and 1:200 for IgG4 specific ELISAs with PBS/10 % (v/v) fetal calf serum, was added and incubated overnight at 4°C. Wells were then incubated with biotinylated mouse anti-human IgE (Pharmingen), followed by streptavidin-biotinylated horseradish peroxide (HRP) complex (Mast Group Ltd.). The assay was developed with o-phenylenediamine substrate solution (Sigma). Plasma samples from 26 uninfected, non-endemic control donors were included in each assay.

For reciprocal inhibition assays, ‘competitor’ recombinant TAL proteins were added at a concentration of 150 μg/ml to a pool of plasma, diluted 1:20 with PBS/10 % (v/v) fetal calf serum, derived from 10 individuals with a known response to each of the respective TAL proteins under investigation. The plasma was then incubated at room temperature for 1 hour before adding to the ‘target’ recombinant TAL protein coated plate and continuing the IgE ELISA described above.

### Statistical analysis

All statistical analysis was conducted in R (version 3.4 (43)). Detection thresholds for IgE seroprevalence were determined by the mean plus 3 standard deviations of non-infected non-endemic control plasma samples (n=26). Egg counts were log transformed prior to analysis (ln(epg + 1)). The significance of binding differences in the cross-reactivity ELISA was tested using sequential t-tests, with p-values adjusted for multiple testing using the Simes-modified Bonferroni correction (44). Specific IgE titers to worm expressed TAL proteins tend to increase following treatment-induced worm death (8). Linear regression analysis was used to investigate associations between 9-week post-treatment SmTAL11-IgE seropositivity alone (results for SmTAL1, 3 and 5 alone are reported elsewhere (3)), and SmTAL1-IgE, SmTAL3-IgE, SmTAL5-IgE and SmTAL11-IgE combined seroprevalence, with intensity of reinfection with *S. mansoni*, after adjustment for age and treatment efficacy (9-week epg). Epidemiological evidence supports the manifestation of processes affecting worm fecundity in *S. haematobium* that do not appear to occur in *S. mansoni* (45). *Schistosoma* parasite load is typically assessed through the measurement of the number excreted eggs; yet, due to the non-linear relationship between CAA and egg excretion in *S. haematobium*, egg counts are an unreliable measure of worm burden, instead the use of CAA is preferable in assessing *S. haematobium* infection intensity (29). In the present study longitudinal follow-up CAA data was not available, so the association between ShTAL1-IgE, ShTAL3-IgE, ShTAL5-IgE and ShTAL11-IgE responses and worm burden was initially analysed using cross-sectional linear models of pre-treatment ShTAL protein IgE seropositivity and baseline CAA with adjustment for age and village of residence. It has been suggested that IgG4 antibodies may compete for the same epitopes and therefore acts to block IgE pathways (46). Where borderline associations were found in regression models, TAL-specific IgG4 was adjusted for to account for any blocking action that specific IgG4 antibodies may have on IgE binding. Logistic regression models were used to explore *S. haematobium* reinfection status at two years post-treatment as a binary outcome, with the justification that detectable egg excretion is an indicator of infection, regardless of worm burden. Post-treatment models were again adjusted for potential confounding predictors. All models were reduced by stepwise removal of non-significant variables. Regression beta coefficients were exponentiated and the results expressed as the geometric mean (GM) ratio.

## Results

### The *S. haematobium* TAL protein family display similar sequence homology to *S. mansoni*

The BLASTp searches resulted in the identification of 13 predicted *S. haematobium* proteins (Table 1). Eleven of these were orthologous to the sequences of SmTAL1–SmTAL5, SmTAL7–SmTAL9 and SmTAL11–SmTAL13, with the EF-hand and DLC domains characteristic of the TAL protein family. A *S. haematobium* protein was also identified with significant orthology to the *S. mansoni* protein SmTAL10. The BLAST searches returned an additional ShTAL (XP_012799749; MS3_0020076) which did not show orthology to existing SmTALs. After BLASTp searches of the *S. mansoni* genome predicted protein dataset, it was shown this ShTAL was an ortholog (87.5% identity) of the previously uncharacterized *S. mansoni* protein assigned here as SmTAL14 (XP_018654160; Smp_146460). Figure 1 shows the alignment of the identified ShTAL proteins by predicted EF hand and DLC domains. Further examination of the Shaem_V3.0 predicted ShTAL5 sequence (XP_035588243) identified a probable error in the annotation of the ShTAL5 and ShTAL6 sequences, merging the two genes. Subsequent alignment of the ShTAL5 Shaem_V3.0 sequences against the Shaem_V1.0 and *S. mansoni* v9 TAL5 and TAL6 sequences shows sequence similarity of TAL5 to the 5’ end and TAL6 to the 3’ end of the Shaem_V3.0 annotation of ShTAL5 (Supporting Information). Furthermore, sequencing of the plasmid from which recombinant ShTAL5 protein was expressed confirmed a coding DNA sequence that aligns to the Shaem_V1.0 annotated sequence with 100% identity (Supporting Information).

**Fig. 1.**
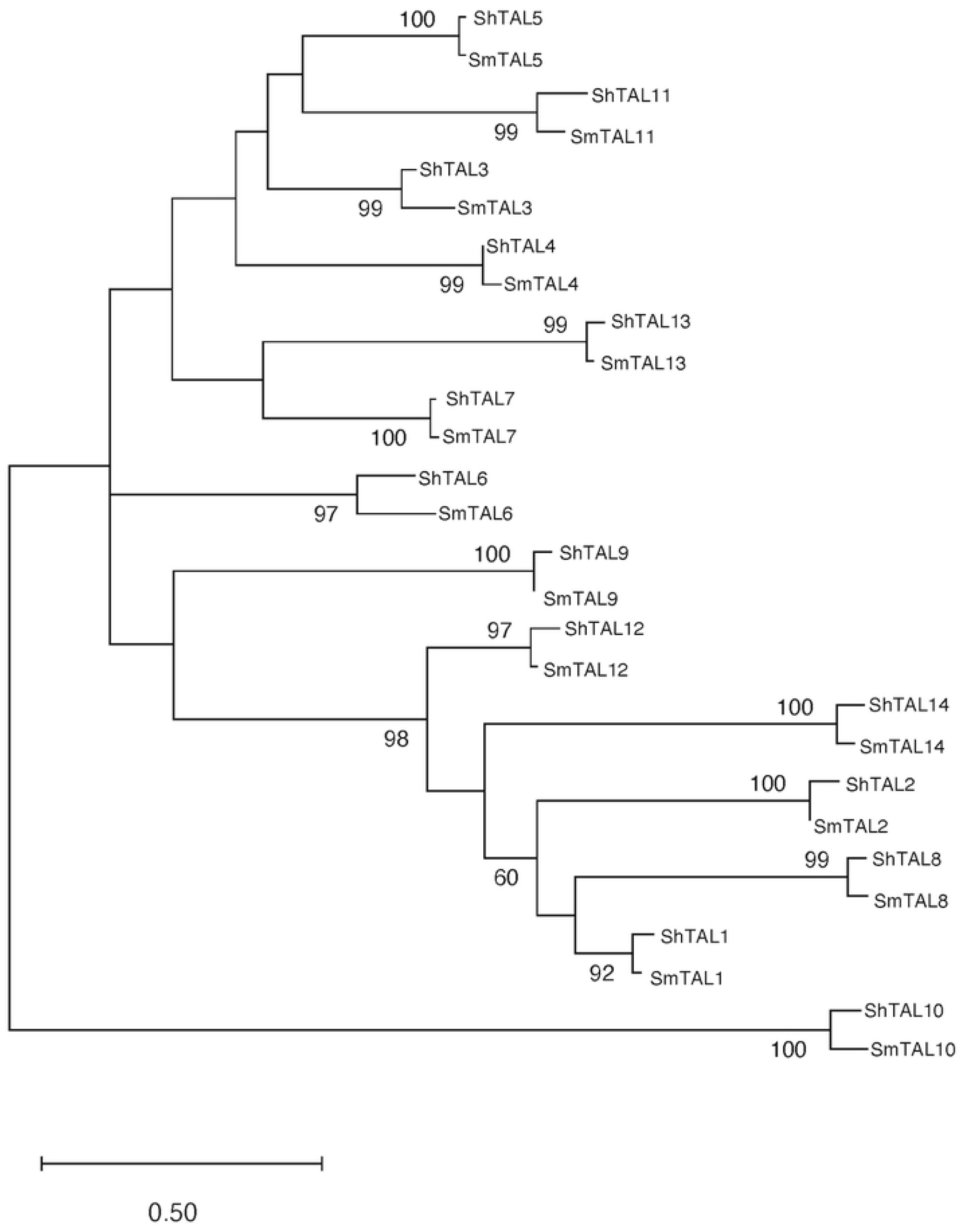
Alignment of amino acid sequences of the *S. mansoni* and likely *S. haematobium* TAL protein family by the predicted TAL protein EF-hand pair (boxed) and predicted DLC region (marked in grey). Conserved Aspartic acid (D) positions within the EF hand domains are indicated in black. * *S. haematobium* Shaem_V1.0 assembly Sequence Performed using MUSCLE (32); accession numbers can be found in Table 1.

**Table 1.**
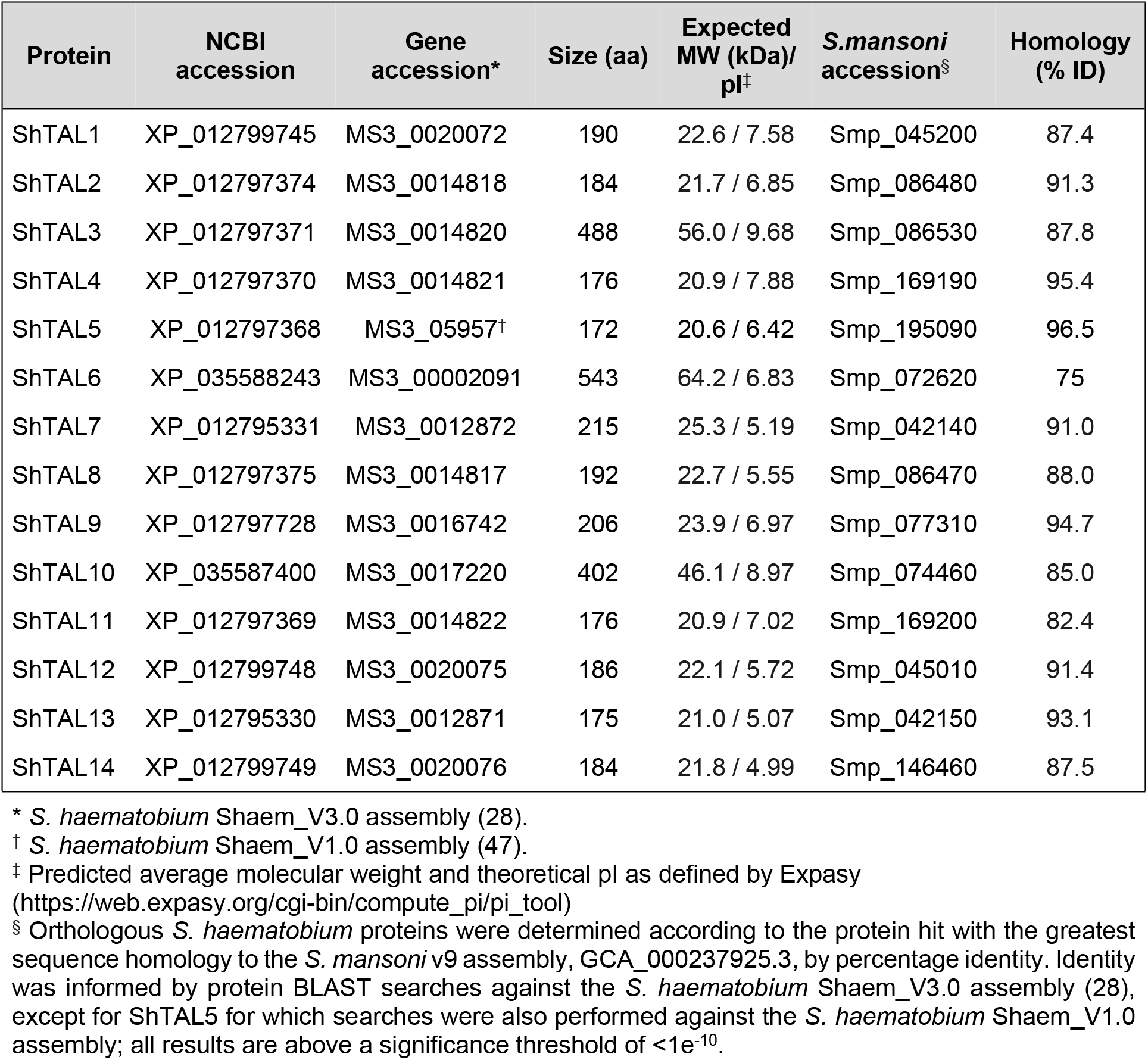
Identity of *S. haematobium* TAL proteins

| Protein | NCBI accession | Gene accession* | Size (aa) | Expected MW (kDa)/ pI<sup>‡</sup> | *S.mansoni* accession<sup>§</sup> | Homology (% ID) |
| --- | --- | --- | --- | --- | --- | --- |
| ShTAL1 | XP_012799745 | MS3_0020072 | 190 | 22.6 / 7.58 | Smp_045200 | 87.4 |
| ShTAL2 | XP_012797374 | MS3_0014818 | 184 | 21.7 / 6.85 | Smp_086480 | 91.3 |
| ShTAL3 | XP_012797371 | MS3_0014820 | 488 | 56.0 / 9.68 | Smp_086530 | 87.8 |
| ShTAL4 | XP_012797370 | MS3_0014821 | 176 | 20.9 / 7.88 | Smp_169190 | 95.4 |
| ShTAL5 | XP_012797368 | MS3_05957<sup>†</sup> | 172 | 20.6 / 6.42 | Smp_195090 | 96.5 |
| ShTAL6 | XP_035588243 | MS3_00002091 | 543 | 64.2 / 6.83 | Smp_072620 | 75 |
| ShTAL7 | XP_012795331 | MS3_0012872 | 215 | 25.3 / 5.19 | Smp_042140 | 91.0 |
| ShTAL8 | XP_012797375 | MS3_0014817 | 192 | 22.7 / 5.55 | Smp_086470 | 88.0 |
| ShTAL9 | XP_012797728 | MS3_0016742 | 206 | 23.9 / 6.97 | Smp_077310 | 94.7 |
| ShTAL10 | XP_035587400 | MS3_0017220 | 402 | 46.1 / 8.97 | Smp_074460 | 85.0 |
| ShTAL11 | XP_012797369 | MS3_0014822 | 176 | 20.9 / 7.02 | Smp_169200 | 82.4 |
| ShTAL12 | XP_012799748 | MS3_0020075 | 186 | 22.1 / 5.72 | Smp_045010 | 91.4 |
| ShTAL13 | XP_012795330 | MS3_0012871 | 175 | 21.0 / 5.07 | Smp_042150 | 93.1 |
| ShTAL14 | XP_012799749 | MS3_0020076 | 184 | 21.8 / 4.99 | Smp_146460 | 87.5 |

\* *S. haematobium* Shaem\_V3.0 assembly (28).
<sup>†</sup> *S. haematobium* Shaem\_V1.0 assembly (47).
<sup>‡</sup> Predicted average molecular weight and theoretical pI as defined by Expasy (https://web.expasy.org/cgi-bin/compute_pi/pi_tool)
<sup>§</sup> Orthologous *S. haematobium* proteins were determined according to the protein hit with the greatest sequence homology to the *S. mansoni* v9 assembly, GCA\_000237925.3, by percentage identity. Identity was informed by protein BLAST searches against the *S. haematobium* Shaem\_V3.0 assembly (28), except for ShTAL5 for which searches were also performed against the *S. haematobium* Shaem\_V1.0 assembly; all results are above a significance threshold of $<1e^{-10}$ .

Amongst the *S. mansoni* and *S. haematobium* TAL orthologues, TAL3, TAL4, TAL5 and TAL11 share greatest sequence identity, with ShTAL 3 and 5 sharing 45.35% sequence identity: ShTAL5 and 11 sharing 46.51% sequence identity, and SmTAL5 sharing 44.77% identity with both SmTAL3 and SmTAL11. Full percentage sequence identity data for this group of TAL proteins can be found in the Supporting Information.

### The TAL proteins can be divided into two principal clades

To identify whether the ShTAL protein phylogenetic topology reflects that of *S. mansoni*, a Maximum Likelihood tree was constructed for the conserved regions detected by Gblocks analysis (Fig. 2). The tree inferred from the refined Gblocks alignment (Log Likelihood −3152.47) shows that the *S. haematobium* proteins are phylogenetically aligned with the *S. mansoni* orthologues. There are two main clades into which the TAL protein family appears to be divided (98% support), one clade including *S. mansoni* and *S. haematobium* TAL1, 2, 8, 12 and 14 proteins, whilst the other includes the remaining *S. mansoni* and *S. haematobium* TAL3, 4, 5, 7, 9, 10, 11, 13 and TAL6 proteins (Fig. 2). Of our immune targets of interest, TAL3, TAL5 and TAL11 therefore cluster in a separate clade from TAL1, the immune target best characterized immuno-epidemiologically for *S. mansoni*, and the only family member characterized immuno-epidemiologically for *S. haematobium*.

**Fig. 2.**
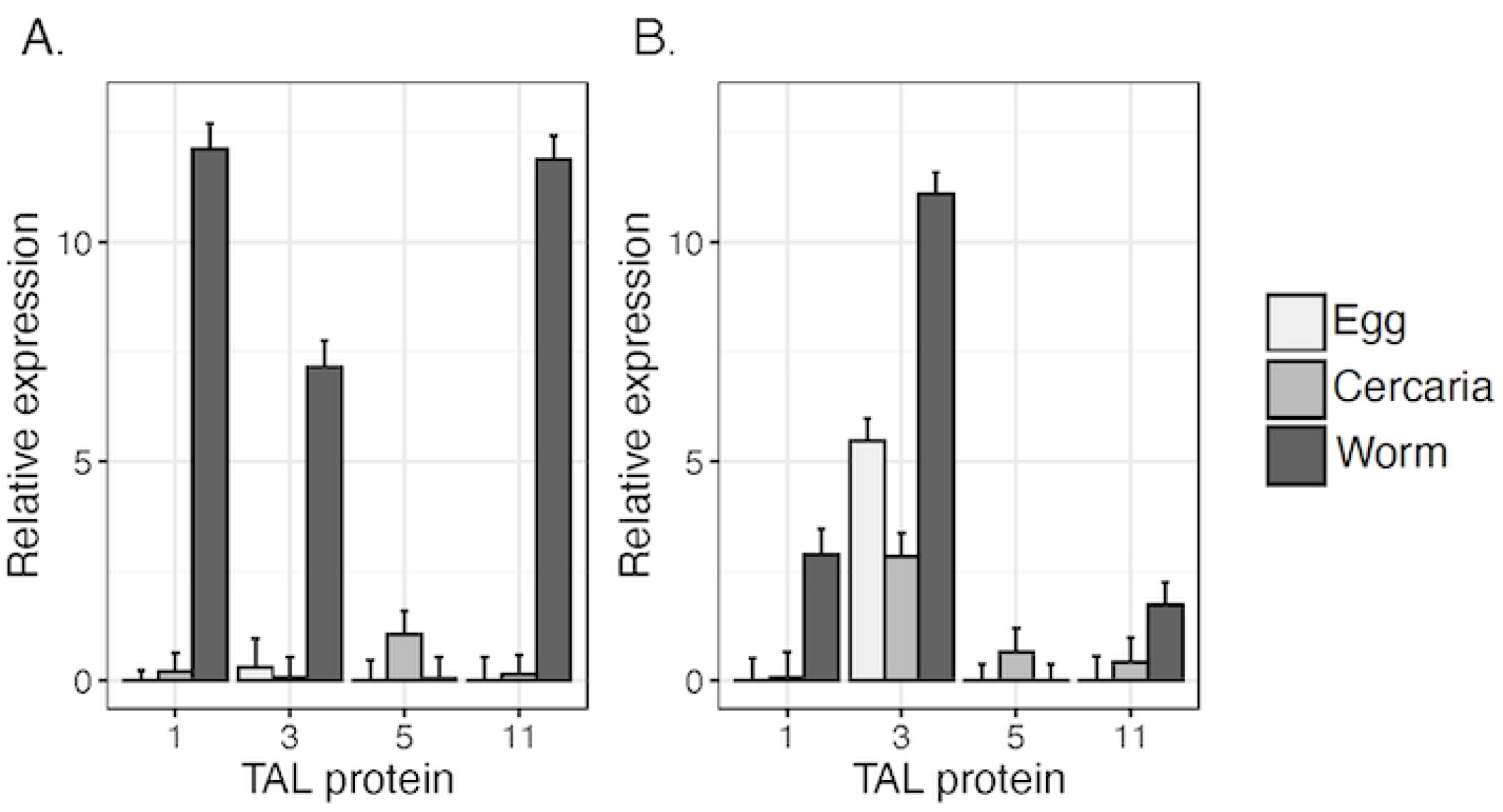
Molecular Phylogenetic analysis by Maximum Likelihood method. Protein sequence analysis of the *S. mansoni* and *S. haematobium* TAL family. The evolutionary history was inferred using the Maximum Likelihood method based on Jones-Taylor-Thornton model for the highly conserved regions detected by Gblocks analysis. The bootstrap consensus tree is drawn to scale, with branch lengths representing the evolutionary distances used to infer the phylogenetic tree (number of substitutions per site). The percentage of replicate trees in which the associated taxa clustered together in the bootstrap test (1000 replicates) are shown next to the branches with a cut-off of 50% and above.

### Transcription profiles of *S. haematobium* TAL proteins vary across life cycle stages and are consistent with a possible adult-worm induced, cercarial response cross-reactive profile

Transcription of *Tal1* is predominant in *S. mansoni* and *S. haematobium* adult worms and absent in the egg and cercarial stages (Fig. 3). Likewise, *Tal3* and *Tal11* are predominantly transcribed in the *S. mansoni* and *S. haematobium* adult worm; though, in contrast to the profile of the *S. mansoni* TAL proteins, *Shtal3* is also transcribed in the egg and cercariae and *Shtal11* is transcribed in the cercariae (Fig. 3). Transcription of *Tal5* is restricted to the cercariae in both species.

**Fig. 3.**
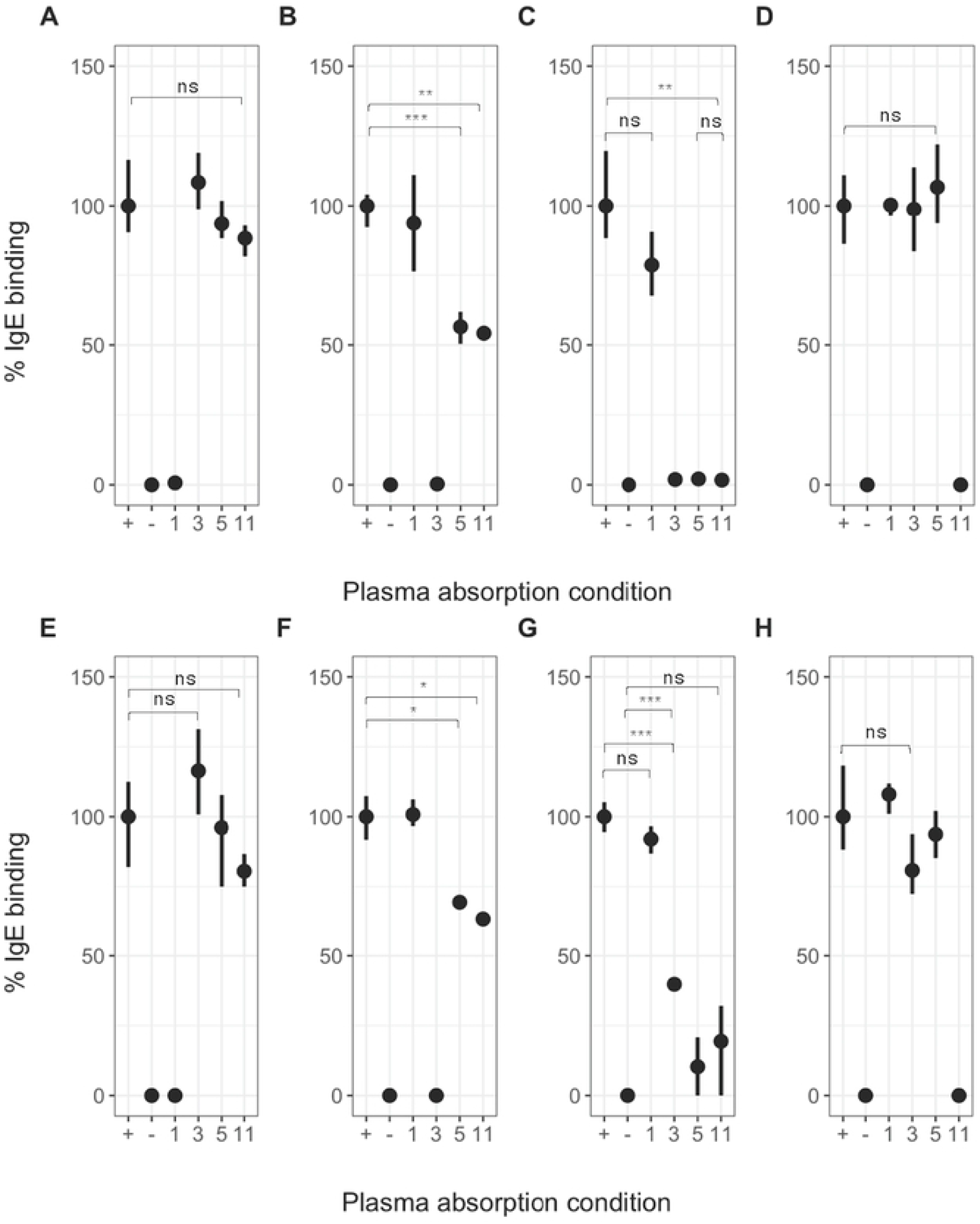
Transcription profiles of *S. mansoni* and *S. haematobium* TAL proteins. Relative lifecycle stage expression of the A) *S. mansoni* and B) *S. haematobium* Tal1, Tal3, Tal5 and Tal11 proteins for egg (light grey), cercaria (medium grey) and adult worm (dark grey). Relative expression was calculated using the delta-delta Ct method (41): Fold gene expression = 2^-(ΔΔCt), where ΔΔCt =ΔCt (Sample) – ΔCt (Control average).

The lifecycle transcription patterns presented here for Shtal3 and Shtal11 concur largely with those reported by Stroehlein *et al*. (28) (Supporting Information), albeit a higher level of transcription found for Shtal3 in the egg in the data shown here.

### IgE specific to TAL11 is cross-reactive with cercarial expressed TAL5 in *S. mansoni* and *S. haematobium*

Despite sequence and lifecycle expression similarities with TAL3, TAL11 has not previously been explored in relation to protective immunity. To elucidate this relationship, the specific post-treatment IgE responses to SmTAL1, 3, 5 and 11 were analysed in relation to their epidemiological profiles (Uganda). IgE responses to the equivalent *S. haematobium* TAL proteins were also examined (Mali). Individuals who were found to have a detectable TAL specific IgE response were identified as an IgE ‘responder’ for a specific TAL (greater than the mean + 3 x standard deviations of the non-endemic control response). 134 individuals (93 % of total Ugandan cohort) had a positive response to one or more of SmTAL1, 3, 5 or 11, with 79 % (11/14) of IgE responders to SmTAL11 also found to be responders to SmTAL1, SmTAL3 and SmTAL5. Likewise, individual IgE responses in the Malian cohort show an overlap in *S. haematobium* TAL protein responses. Sixty-five percent (28/43) of ShTAL11 responders were quadruple responders, with IgE also measurable for ShTAL1, ShTAL3 and ShTAL5. Indicating that individuals seropositive for TAL11-IgE may be a subset of those with a detectable TAL5, TAL3 and TAL1-specific IgE response in *S. mansoni* and *S. haematobium* infection.

The overlap in individual responses to multiple TAL proteins indicates cross-reactivity could be occurring. To ascertain whether IgE to SmTAL11 and the equivalent *S. haematobium* antigens demonstrate a similar cross-reactivity profile to that previously shown between SmTAL3 and SmTAL5, reciprocal inhibition ELISAs were conducted (Fig. 4).

**Fig. 4.**
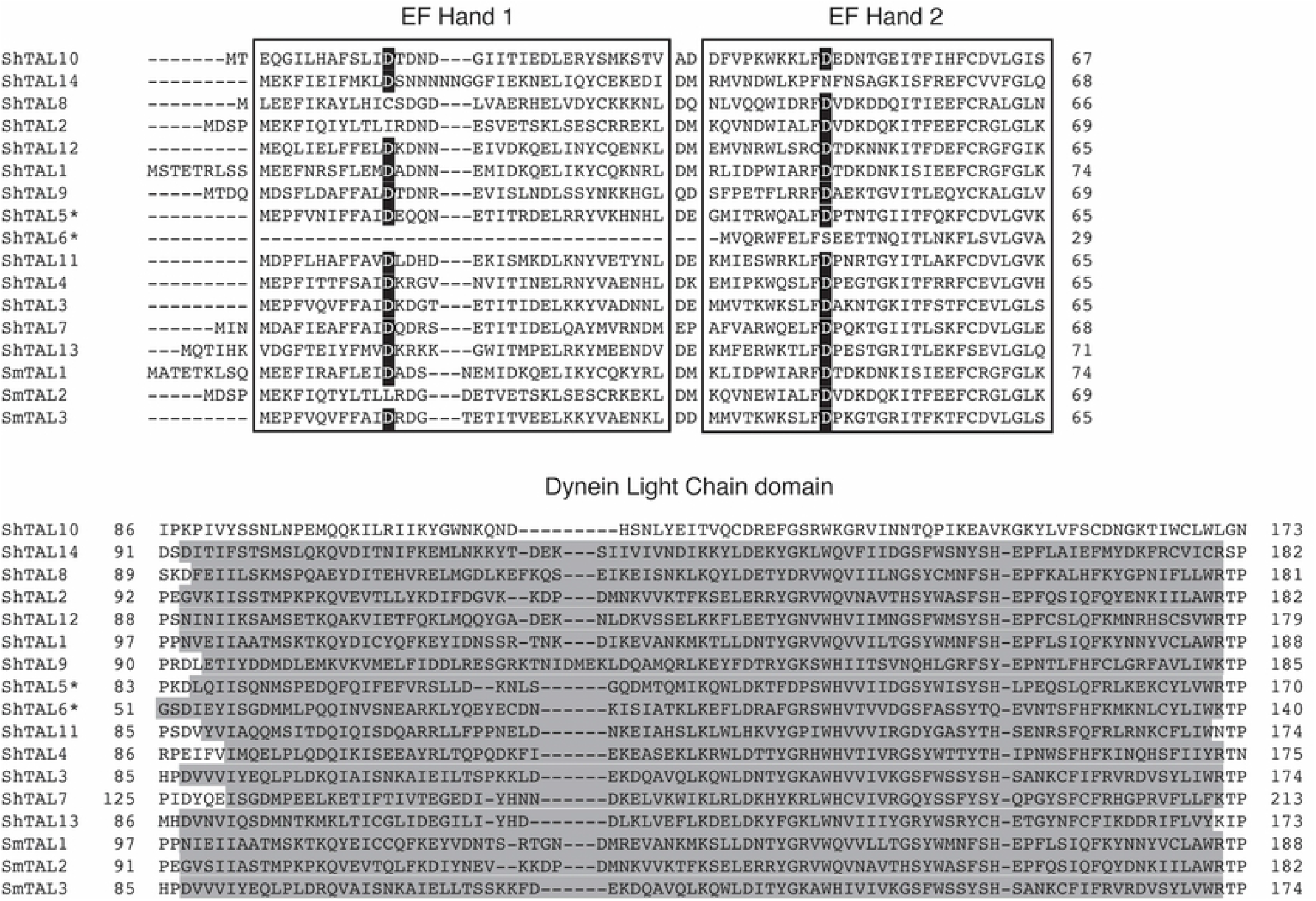
Cross-reactivity of TAL-IgE responses in *S. mansoni* and *S. haematobium*. Reciprocal inhibition ELISA using pooled plasma (n=10) from individuals with demonstrable IgE response to Sm or Sh TAL1, 3, 5, and 11. IgE binding to solid-phase SmTAL1 (A), SmTAL3 (B), SmTAL5 (C), SmTAL11 (D) and ShTAL1 (E), ShTAL3 (F), ShTAL5 (G) and ShTAL11 (H) was inhibited by pre-incubation with antigens Sm or Sh TAL1, 3, 5 or 11, respectively, in solution at 150 μg/ml. Positive control plasma was not pre-incubated with antigen. Negative control contained no plasma. Samples were run in triplicate; points indicate mean percentage IgE concentration relative to positive control. Error bars indicate minimum & maximum value. The significance of binding differences was tested using t-test, with p-values adjusted for multiple testing using the Simes-modified Bonferroni correction (38), where *** p < 0.001, ** p < 0.01, * p < 0.05 and ns = non-significant.

Complete inhibition of IgE binding to SmTAL5 following pre-incubation with either SmTAL3 (t = 0.756; p = 0.493) or SmTAL11 (t = 1.225; p = 0.288), comparable to pre-incubation with SmTAL5 (Fig. 4C), was observed. In contrast, although pre-incubation of plasma with ShTAL3 did significantly suppress IgE binding to ShTAL5 (t = −31.956; p = 0.001, compared to negative control) in the *S. haematobium* assay, binding was not completely suppressed, instead resulting in an approximately 60 % reduction in the IgE signal from the positive control (Fig. 4G); whereas the level of inhibition following pre-incubation with ShTAL11 was not significantly different from pre-incubation with either ShTAL5 (t = −0.794; p = 0.480) or the negative control (t = −1.970; p = 0.188) across the replicates (Fig. 4G), indicating full suppression of the binding of IgE to ShTAL5 by ShTAL11. Binding of plasma IgE to solid-phase SmTAL3 was reduced by almost 50 % following pre-incubation of the plasma with both SmTAL5 (t = 8.497; p = 0.001) and SmTAL11 (t = 11.589; p = 0.006) (Fig. 4B). Suppression of ShTAL3 binding was also significantly inhibited by pre-incubation with the equivalent *S. haematobium* recombinant proteins compared to the positive control (Fig. 4F, ShTAL5 [t = 6.667; p = 0.021] and ShTAL11 [t = 7.987; p = 0.015]). In contrast, IgE binding to solid-phase SmTAL11 was not significantly inhibited by preincubating plasma with SmTAL1, SmTAL3 or SmTAL5 (Fig. 4D); nor was a reduction in ShTAL11 seen for serum pre-incubated with the equivalent ShTAL proteins (ShTAL1 (t = −0.809; p = 0.487), ShTAL3 (t = 1.684; p = 0.175) or ShTAL5 (t = 0.598; p = 0.592), all compared to positive control) (Fig. 4H). Finally, the assay did not support cross-reactivity between TAL1 and other members of the TAL protein family in *S. mansoni* (Fig. 4A) or *S. haematobium* (Fig. 4E). Binding of plasma IgE to each solid-phase, plate-bound TAL protein was completely inhibited by pre-incubation of the plasma with the respective soluble TAL antigen, as compared to the negative control, indicating that all available antigen binding sites, or paratopes, had been blocked by ‘free’ antigen in solution. The results of the cross-reactivity assay therefore indicate that SmTAL11 and ShTAL11 are also cross-reactive components of the IgE response to the cercarial expressed TAL5 but that this cross-reactivity is one-directional, with no reciprocal inhibition of response observed.

### SmTAL11 IgE is associated with reduced reinfection

Population based linear regression models demonstrate that increasing host age and having a positive IgE response to SmTAL11 was significantly associated with reduced intensity of reinfection at two years post-treatment (GM ratio: 0.18; p < 0.01). Seropositivity for combined SmTAL1, SmTAL3, SmTAL5 and SmTAL11 specific IgE was also significantly associated with reduced reinfection intensity at two years post-treatment, even after accounting for age in the model (Table 2). Pairwise interaction terms between the independent variables were not significant and were therefore removed from the models. These results lend further support to the hypothesis that SmTAL11 is associated with the development of delayed concomitant immunity.

**Table 2.** Geometric mean odds ratios describing the association between post-treatment IgE responses to SmTAL antigens and intensity of reinfection at two years post-treatment.

|  | SmTAL1, 3, 5, 11 |  |
| --- | --- | --- |
|  | GM ratio (95% CI) | p-value |
| Age (years)<sup>†</sup> |  |  |
| 10 to 14 | 0.96 (0.32–2.88) | ns |
| 15 to 24 | 0.38 (0.11–1.28) | ns |
| 25 to 34 | 0.17 (0.06–0.52) | ** |
| 35 to 60 | 0.10 (0.03–0.29) | *** |
| 9-week egg count<sup>‡</sup> | 1.14 (0.89–1.46) | ns |
| SmTAL seropositivity | 0.19 (0.04–0.87) | * |

\*\*\* $p < 0.001$ , \*\* $p < 0.01$ , \* $p < 0.05$ , ns: non-significant; <sup>†</sup> values compared to 7- to 9-year-old age group; <sup>‡</sup> proxy for treatment efficacy. Models are adjusted for age and 9-week egg count, as a measure of treatment efficacy.

### *S. haematobium* TAL-specific IgE is associated with lower infection intensity both pre- and posttreatment

Being seropositive for ShTAL3-IgE (GM ratio: 0.48; 95 % CI: 0.25–0.94; p = 0.03) was negatively associated with baseline infection intensity in cross-sectional models of anti-ShTAL responses and infection intensity (as measured by CAA), that accounted for age, village of residence and the relevant TAL-specific IgG4 response. ShTAL5-IgE (GM ratio: 0.79; 95 % CI: 0.40–1.57; p = 0.50) also demonstrated a negative association with baseline infection intensity, though not significantly. Specific IgG4 responses were significantly associated with higher baseline CAA (ShTAL3-IgG4: GM ratio: 4.56; 95 % CI: 2.26–9.17; p < 0.0001; ShTAL5-IgG4: GM ratio: 3.95; 95 % CI: 2.04–7.64; p = 0.0001). Adjusting for ShTAL3-specific IgG4 was required to maintain the effect of the negative association between seropositivity for ShTAL3-IgE and baseline infection. The results of models without adjustment for TAL-IgG4 are provided in the Supporting Information Table S2. When combined TAL responsiveness was examined, no significant associations were seen for double (ShTAL1 and 3), triple (ShTAL1, 3 and 5), or quadruple (ShTAL1, 3, 5 and 11) responders (Supporting Information Table S3). Post-treatment increases were observed in specific IgE antibody titers to ShTAL1 (t = −3.257, p = 0.001), ShTAL3 (t = −4.108, p < 0.0001), and ShTAL5 (t = −2.0055, p = 0.05) and ShTAL11 (t = −3.8045, p = 0.0002).

Although two-year follow-up CAA data was not available for analysis of the association between post-treatment ShTAL-IgE seropositivity and intensity of reinfection, logistic regression models were built to test whether post-treatment ShTAL protein IgE seropositivity was associated with reinfection status as determined by detection of eggs in the urine at two years. Being IgE seropositive to ShTAL1, 3 and 5 (GM ratio: 0.39; 95 % CI: 0.18–0.84; p = 0.02) was significantly associated with protection against reinfection and remained so when age and village were accounted for in the model (Table 3; GM ratio: 0.29; 95 % CI: 0.09 – 0.90; p = 0.03). Sex, treatment efficacy and TAL-specific IgG4 were also considered as confounding factors, but these were found to be non-significant and were removed from the model. Being a quadruple responder to ShTAL1, 3, 5, and 11 was non-significant (Table 3).

**Table 3.** Association between 9-week post-treatment ShTAL-specific IgE responses and reinfection status at two years. Models adjusted for age and village. Results are displayed for total cohort (n = 174).

|  | ShTAL1, 3, 5 |  | ShTAL1, 3, 5, 11 |  |
| --- | --- | --- | --- | --- |
|  | GM ratio (95% CI) | P-value | GM ratio (95% CI) | P-value |
| Age (years) <sup>†</sup> |  |  |  |  |
| 7–9 | 1.49 (0.33–7.96) | ns | 1.44 (0.32–7.55) | ns |
| 10–14 | 0.43 (0.10–1.89) | ns | 0.36 (0.08–1.53) | ns |
| 15 – 24 | 0.03 (0.01–0.15) | *** | 0.03 (0.01–0.13) | *** |
| 25 – 29 | 0.06 (0.01–0.22) | *** | 0.04 (0.01–0.16) | *** |
| 30+ | 0.03 (0.01–0.12) | *** | 0.03 (0.01–0.12) | *** |
| Village<sup>‡</sup> | 27.05 (8.45–107.53) | *** | 23.33 (7.57–88.05) | *** |
| TALx-IgE | 0.29 (0.09–0.90) | * | 0.95 (0.17–5.28) | ns |

\*\*\*p<0.001, \*p<0.05, ns non-significant;
<sup>†</sup> values compared to 5- to 6-year-old age group;
<sup>‡</sup> values provided for high compared to moderate transmission intensity villages

## Discussion

The IgE antibody response has long been associated with both allergy and immunity in parasitic infections, with sequence identity and structural similarity between environmental allergens and parasite proteins having been utilized in the prediction of allergen-like protein families in *S. mansoni* (20). These structural similarities can give rise to cross-reactive host antibody responses upon exposure to allergens (48). Immunogenic cross-reactivity has been observed between several parasite allergen-like proteins (26). Cross-reactivity between malaria variant surface antigens has also been identified as a mechanism to evade the host immune response and drive disease chronicity (49). The importance of inter-species cross-reactivity and boosting of the host immune response to similar proteins has also been recognized in the identification of potential transmission blocking vaccine targets (50). In human schistosomiasis, cross-reactivity has been observed between Venom Allergen-Like (VAL) proteins in *S. mansoni*, a family of secreted proteins identified as a potential intervention target (51, 26).

These studies establish processes of cross-reactivity as vital in our understanding of host-pathogen interactions; interactions that can lead to exacerbation of pathology, or aid or hinder the development of protective responses, key knowledge for assessing therapeutic targets. Here, for the first time, we conducted comparative analysis of the Tegument Allergen Like (TAL) protein family across two of the major human infecting schistosomes, with a particular focus on their role at the host-parasite interface, humoral cross-reactivity, and the resultant host protection.

We identified 14 TAL proteins in the *S. haematobium* V3.0 genome, including an orthologue of a previously uncharacterized *S. mansoni* protein, assigned TAL14. Analogous with the phylogeny of *S. mansoni* TAL proteins, the *S. haematobium* TAL phylogeny revealed that the TAL proteins cluster within two subclades (98% support): the Group 1 TAL proteins comprising TAL1, 2, 8, 12 and 14 and the Group 2 proteins comprising the remaining TAL proteins: TAL3, 4, 5, 6, 7, 9, 10, 11, and 13. Amongst the Group 2 proteins TAL3, 4, 5 and 11 share the greatest sequence identity with *S. mansoni* and *S. haematobium* TAL3, 5 and 11 sharing moderate sequence identity, between 30–60 %. Whilst Aalberse (52) noted that shared sequence identity greater than 50% is likely to result in cross reactivity, cross reactivity between antibodies against SmVAL proteins sharing at least 30% sequence identity has been demonstrated (26). Fitzsimmons *et al*. (3) have previously evidenced cross-reactivity between SmTAL3 and SmTAL5, conferring IgE-mediated partially protective immunity, yet the closely related family member TAL11 has not previously been characterized immuno-epidemiologically for either parasite species.

Identifying the developmental expression profiles for the ShTAL proteins is vital in understanding the human immune response to this family of proteins. Each lifecycle stage evokes a distinct immune response as a result of the anatomical environment in which they reside, and the frequency of antigen exposure to the immune system (3,8). Several parasites employ this complex biology of stage specific antigenic expression as a mechanism to evade detection by the human host (53). In *S. mansoni, Smtal5* is expressed throughout the cercaria (8) and single cell transcript analysis indicates that it is also expressed in the transforming schistosomula (54), evidence for its potential as a key target of the protective response directed against these parasite life stages. In examining the developmental expression profiles of the ShTAL proteins proposed to be associated with protective human IgE responses, we found that ShTAL proteins are also developmentally transcribed. Although the expression profile of *S. haematobium* TALs does differ slightly from that of *S. mansoni*, the lifecycle stage expression of *Shtal1, 3, 5* and *11* remains consistent with the hypothesis that antigen specific IgE induced by worm death and cross-reactivity to ShTAL5 drives protective immunity in *S. haematobium*. This analysis is supported by the transcription data published alongside the most recent *S. haematobium* genome annotation (28). In addition, post-treatment increases observed in specific IgE antibody titers to ShTAL1, ShTAL3 and ShTAL11 are in accordance with the patterns seen in *S. mansoni* (8). This corresponds with predominant expression of these proteins in the adult worm, and their exposure to the host immune system through drug induced worm death.

Secreted and surface expressed proteins have garnered significant interest in the search for vaccine targets. The schistosome tegument plays an important role at the host-parasite interface and, whilst it associated with immune evasion strategies, it is also considered to be the parasite structure most susceptible to immune-mediated attack by the host (55). Several surface-expressed schistosome proteins that induce strong specific IgG isotype responses are differentially expressed throughout the parasite lifecycle and are subsequently under investigation in the drive to identify suitable vaccine candidates (56–58). Conversely the TAL proteins evoke a strong IgE-mediated protective response and, whilst an IgE-mediated immune response does not provide an ideal basis for vaccine development (6), TAL proteins share several characteristics with the most promising protein candidates in the development of protective technologies against schistosome transmission. It is, however, the association with worm death (either natural or drug induced) that makes examination of the host’s immune response to the TAL protein family so critical, since mass drug administration underpins schistosomiasis control. Understanding interactions at the host/parasite interface enable better assessment of the impact these dynamics may have on schistosome transmission and subsequent policy decision-making going forward.

IgE mediated protective immunity to *S. mansoni* has previously been proposed to result from SmTAL3-specific IgE cross-reactive to SmTAL5 that is exposed to the human host on the invading cercaria and early schistosomula (3). SmTAL5 is expressed at comparatively low levels, hence it is thought that antibodies may not be generated to SmTAL5 in sufficient levels to be efficacious. In the analysis presented here we find that a similar cross-reactive relationship to that previous observed between SmTAL3 and SmTAL5 is seen between SmTAL5 and SmTAL11, with pre-incubation with SmTAL11 resulting in complete inhibition of IgE binding to SmTAL5. Preincubation with SmTAL5 and SmTAL11 also reduces SmTAL3 binding; yet pre-incubation with SmTAL3 and SmTAL5 has no effect on SmTAL11 binding. This suggests that either SmTAL3 is not the sole cross-reactive antigen responsible for the development of partially protective immunity and that SmTAL11 may also share IgE epitopes with SmTAL5, or that the cross-reactive epitope is shared by all three closely related TAL proteins. The lack of inhibition of SmTAL11 binding indicates that specific IgE may bind the SmTAL11 epitope with higher affinity compared to the other TAL proteins, resulting from disassociation from the competing antigen upon exposure to the solid phase TAL11. Furthermore, regression models found the *S. mansoni* TAL11-specific IgE response to be strongly associated with reduced reinfection intensity at two years. This association was also seen for those with a positive response to multiple SmTAL proteins (SmTAL1, 3, 5 and 11), which was expected due to the degree of overlap between SmTAL-IgE responders.

Antibody affinity studies would improve our understanding of this relationship still further; however, there are several challenges regarding experimental design and the nature of specific antibody responses in serum from schistosomiasis endemic areas, most notably the polyclonal nature of the antibodies and the presence of non-IgE antibodies that are specific to the same antigens.

*S. haematobium* demonstrates similar cross-reactivity profiles between ShTAL3, ShTAL5 and ShTAL11 to those seen for *S. mansoni* proteins, although a few key differences between the responses are observed. The extent to which ShTAL3 binding is inhibited by ShTAL5 and ShTAL11 is not as high in *S. haematobium* compared to the equivalent *S. mansoni* TAL protein inhibition. Furthermore, preincubation with ShTAL3 does not appear to completely inhibit ShTAL5 binding. Unlike the equivalent *S. mansoni* protein, the transcription data presented here suggest that ShTAL3 has a high level of expression in the egg, in addition to expression in the adult worm. ShTAL3-specific IgG4 responses may consequently block or regulate the cross reactive IgE response, to reduce the pathology that would result from high IgE responses to tissue trapped eggs (59).

In the study population-based analysis, a protective effect of IgE to ShTAL proteins was observed, with having a detectable IgE response to a combination of ShTAL1, ShTAL3 and ShTAL5 significantly associated with a lack of reinfection at two years, providing evidence for the progressive development of a protective IgE response, similar to that demonstrated in *S. mansoni*. Regression models did not, however, lend support to the contribution of *S. haematobium* TAL11 to partially protective immunity. The power to detect an association between ShTAL11 seropositivity and reduced reinfection, however, was limited in this study by the number of individuals seropositive for ShTAL11 specific IgE. Furthermore, CAA titers at reinfection may be more sensitive at detecting this relationship due to a closer reflection of true reinfection rates (60) that are not confounded by a possible anti-fecundity immune response (29, 45). Protective cross-reactivity between ShTAL11 and ShTAL5 cannot therefore be excluded by this study, particularly in view of our inhibition assay findings, in which ShTAL11 had greater binding to ShTAL5 than ShTAL3.

This study utilized immuno-epidemiological inquiry, in combination with *in silico* and parasite transcriptional analyses to explore the relationship between the TAL protein family expression and human immunity to reinfection. It provides important development in our understanding of protective immune responses to *S. mansoni* and for the first time, comparative analysis of this important protein family for *S. haematobium*, the most prevalent of human infecting schistosomes. We provide evidence for a cross-reactive delayed concomitant immunity driven protective response in *S. haematobium*, in line with previous observations for *S. mansoni*. In addition, overlapping IgE seroprevalence to multiple TAL proteins, combined with observable cross-reactivity between TAL protein family members, suggests that adult worm expressed SmTAL11 and ShTAL11 may also drive development of the protective IgE immune response, thus providing further support to the hypothesis of delayed concomitant immunity to these parasites of great public health importance.

## Notes

### Author contributions

RCO manuscript authorship and preparation, bioinformatic analysis, cloning, protein production, serology and statistical analysis; HAD qPCR experiments and analysis, cloning and protein production; CMF cloning and protein production; MS and EMT coordination of field data collection; IWC bioinformatic analysis and manuscript preparation, and SW project conception, manuscript preparation.

